## Supplementary material for "X-Ray Structure and enzymatic study of a Bacterial NADPH oxidase highlight the activation mechanism of eukaryotic NOX": PetitVermot et al 2023-SI revised

**to**

[∂]ALPX S.A.S. 71, Avenue des Martyrs, Grenoble, France

[£] CNRS-Lyon 1 University Laboratory 5086, IBCP, Lyon, France.

[f] Institut Universitaire de France (IUF), Paris, France

1. *These two authors, listed in alphabetical order, contributed equally to the work.*

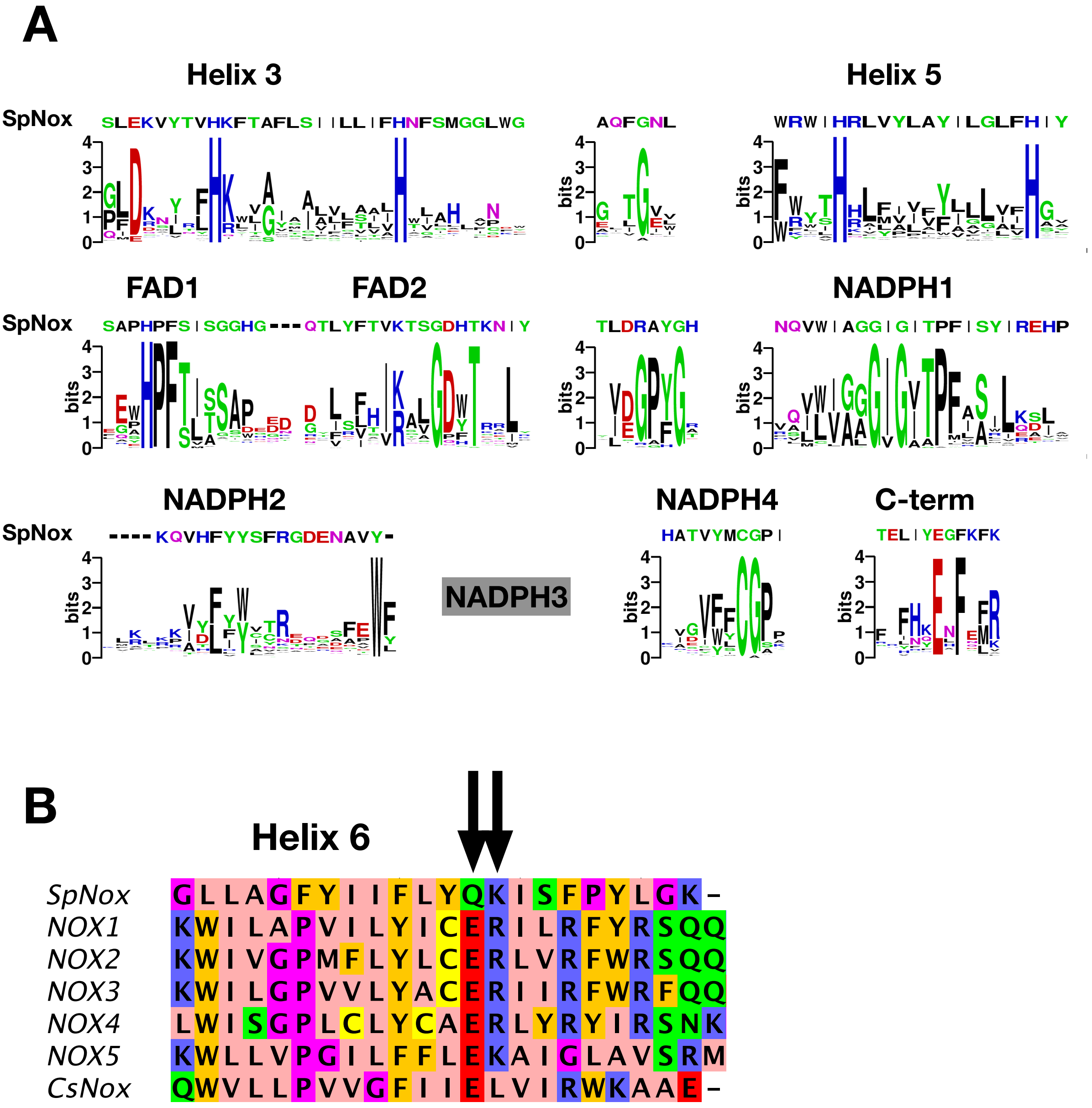


**Fig S1: NOX-specific sites conserved in SpNOX.** A). Alignment of the SpNOX sequence with motifs identified in eukaryotic NOX by Massari et al (61). Alignments of TM and DH domains including prokaryotic and eukaryotic NOX sequences, as described in Materials and Methods, were used to construct logos of subsequences identified as NOX-specific sites in (61). B). The region of human NOX1-5, CsNOX, and SpNOX spanning TM helix 6 and into the first beta strand of the DH domain was aligned to identify the putative end of the TM domain and beginning of the DH domain. Arrows indicate the ER ($E_{283}R_{284}$in NOX2 numbering) pair highly conserved in eukaryotic NOX, and the corresponding $Q_{180}K_{181}$ pair in SpNOX.

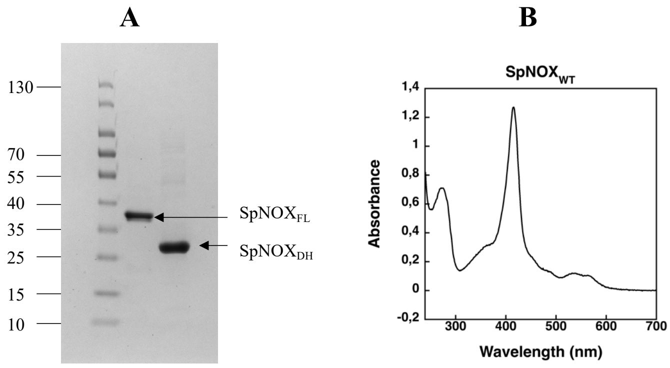


**Fig S2: SDS PAGE of purified SpNOX and SpNOX$_{DH}$.** A). SpNOX full length (FL) and SpNOX DH separated by SDS–PAGE. B). UV-visible spectrum of SpNOX at 5.8 µM (heme concentration) in 50 mM Tris-HCl, pH7, 300 mM NaCl, 0.0025% LMNG.

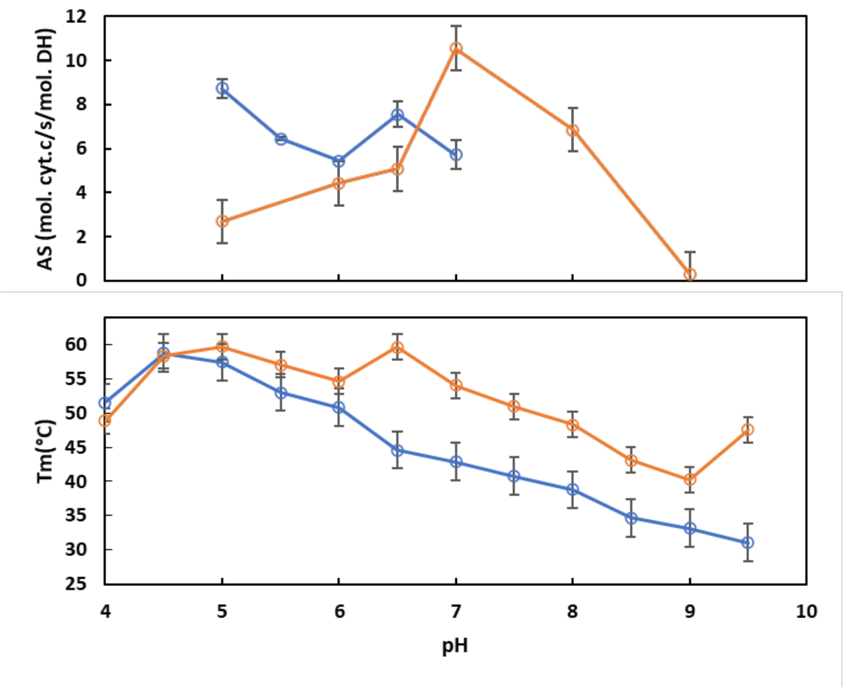


**Fig. S3: Comparison of pH dependence of SpNOX and SpNOX$_{DH}$ thermostability and activity.** Experiments were conducted in buffers at different pH at 0.1M, 300 mM NaCl, and 5% glycerol (see materials & methods). A). Dependence of cytochrome *c* reductase activity of SpNox$_{DH}$ (–) and SpNox (–) on pH. B). Dependence of SpNOX (–) and SpNOX (–) thermostability on pH.

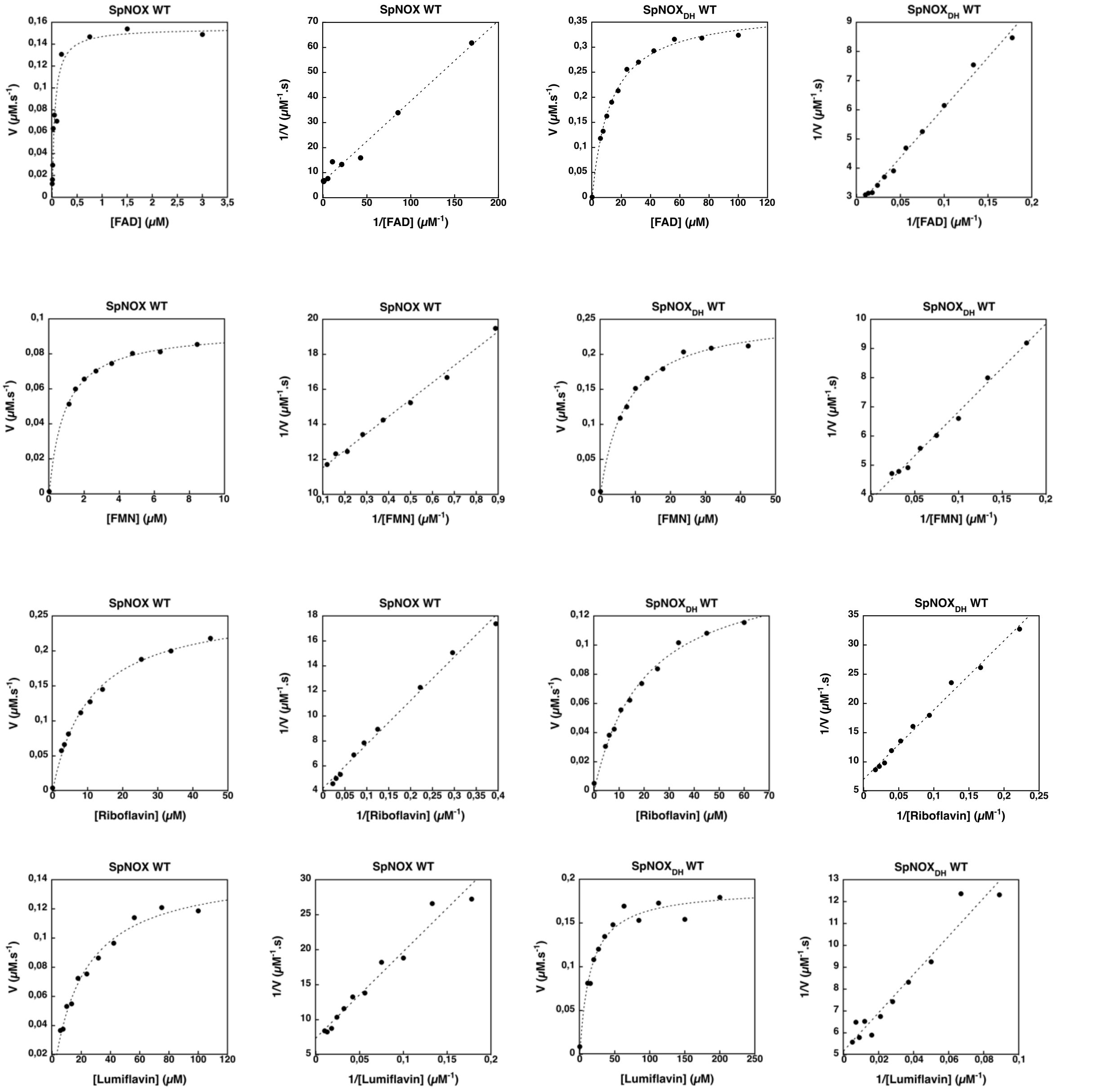


**Fig. S4: Michaelis Menten analysis of SpNOX and SpNOX$_{DH}$ as a function of the flavin substrate**. Cytochrome *c* reductase activity was monitored at 550 nm. For all flavin substrates the Michaelis Menten plots and the Lineweaver-Burk plots are presented. $K_m$ and $k_{cat}$ extracted from these experiments are presented in Table 2.

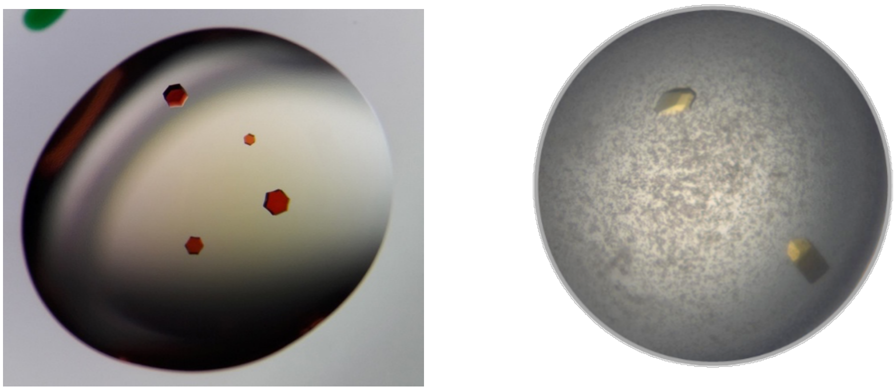

**Fig S5: Crystals of SpNOX F397W (left) and SpNOX$_{DH}$ F397W (right).**

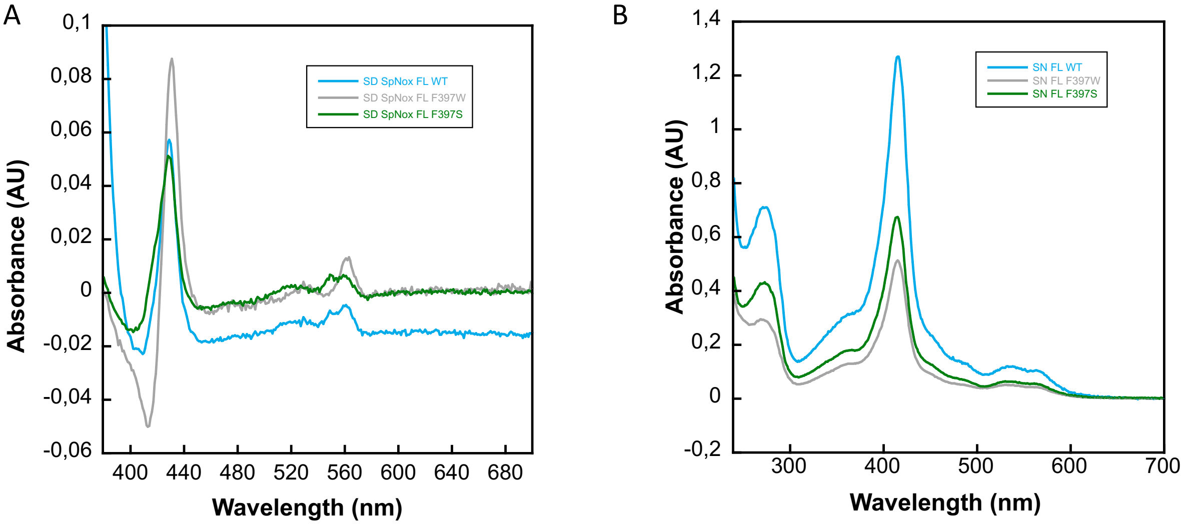


**Fig. S6: Characterization of SpNOX, SpNOX-F397W and SpNOX-F397S mutants*:*** A). Difference spectra (reduced-oxidized) of membrane preparations from *E.coli* expressing SpNox FL WT (blue), SpNox FL F397W (grey) and SpNox FL F397S (green) mutants verify the presence of functional hemes. B). UV-visible spectral characterization of purified SpNox FL WT, SpNox FL F397W and SpNox FL F397S in 50 mM Tris-HCl, pH7, 300 mM NaCl, 0.0025% LMNG (colors as in A). Spectra are not normalized with concentration.

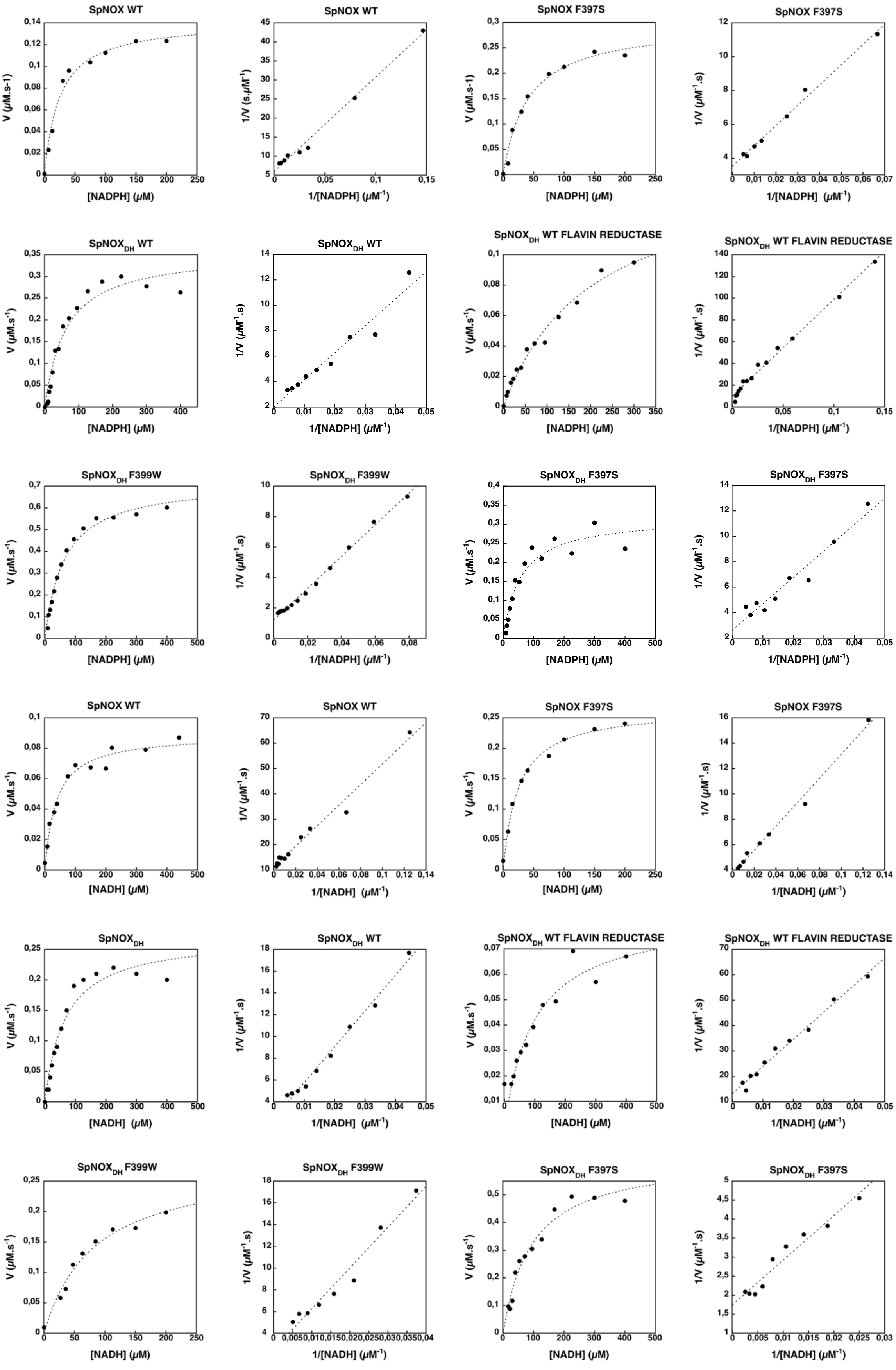


**Fig. S7: Michaelis Menten analysis of wild type and mutant SpNOX and $SpNOX_{DH}$ as a function of the nicotinamide-based electron donor**. Cytochrome *c* reductase activity was monitored at 550 nm, and, in panels so labeled, flavin reductase activity was monitored at 340

nm. For all flavin substrates the Michaelis Menten plots and the Lineweaver-Burk plots are presented. $K_m$ and $k_{cat}$ extracted from these experiments are presented in Table 3.

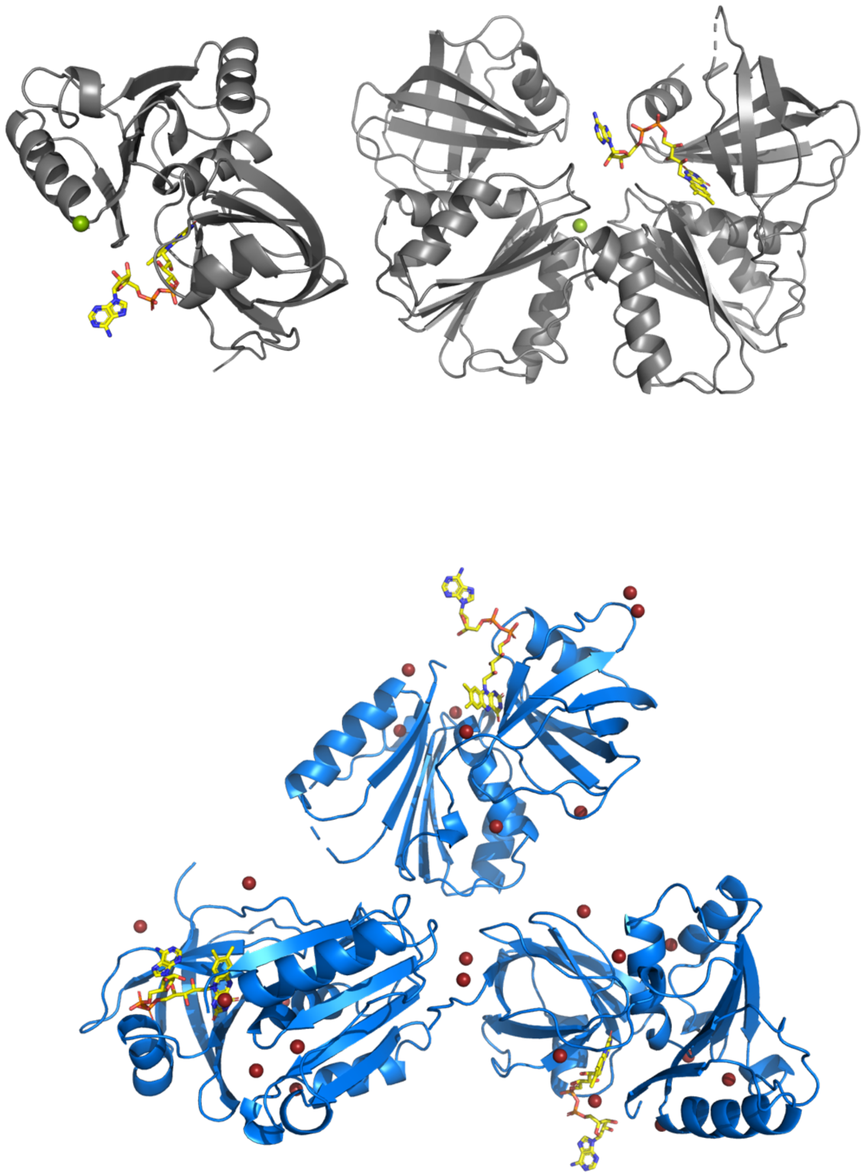

**Figure S8: Asymmetric unit of crystal of SpNOX$_{DH}$ WT (top) and of SpNOX$_{DH}$ F397W (bottom).** FAD is depicted as sticks, colored by atom. Chloride ions are shown as green spheres, bromide ions used for SAD data acquisition as red spheres.

| Data collection | | | |
|---|---|---|---|
| | **DH F397W** | **DH WT** | **FL F397W** |
| Space group | $P4_12_12$ | $P4_12_12$ | $P6_422$ |
| Cell *a*, *b*, *c* (Å) | 104.62, 104.62, 142.68 | 104.88, 104.88, 139.29 | 145.97, 145.97, 153.62 |
| Angles α, β, γ (°) | 90, 90, 90 | 90, 90, 90 | 90, 90, 120 |
| Resolution (Å) | 58.94-1.94 (1.97-1.94) | 46.9-2.50 (2.60-2.50) | 47.82-3.62 (3.95-3.62) |
| $R_{merge}$ | 0.117 (1.829) | 0.140 (2.441) | 0.072 (2.1) |
| $I/\sigma$ | 21.2 (2.1) | 18.99 (1.46) | 19.6 (1.7) |
| Completeness (%) | 98.4 (95.6) | 100.0 (100.0) | 47.9 (10.7)<br>90.9* (85.4*) |
| Redundancy | 13.8 (12.9) | 25.96 (26.86) | 20.4 (20.1) |
| Ellipsoid | n/a | n/a | 0.894 a*-0.447 b*, b*, c* |
| **Refinement** | | | |
| Resolution (Å) | 36.99-1.94 | 46.9-2.5 | 47.82-3.62 |
| Reflections / free % | 57709 / 5.00 | 27532 / 5.00 | 11510 / 4.45 |
| $R_{work}$ / $R_{free}$ | 0.190 / 0.225 | 0.197 / 0.290 | 0.262 / 0.320 |
| B-average (Å$^2$) | 38.64 | 70.2 | 201.0 |
| R.m.s.d<br>Bond lengths (Å)<br>Bond angles (°) | <br>0.012<br>1.55 | <br>0.007<br>1.456 | <br>0.0071<br>1.6931 |
| Ramachandran (%)<br>Favored<br>Outliers | <br>98.6<br>0.0 | <br>93.7<br>0.0 | <br>87.1<br>1.0 |

**Table S1. X-ray data collection and refinement statistics of SpNOX$_{DH}$ WT, SpNOX$_{DH}$ F397W and full length SpNOX.**

*Highest resolution shell is shown in parenthesis.

Data has been fitted to the ellipsoid defined by the following parameters:

Diffraction limits & principal axes of ellipsoid fitted to diffraction cut-off surface:

5.308 Å, 1.0000 0.0000 0.0000 0.894 a*-0.447 b*

5.308 Å, 0.0000 1.0000 0.0000 b*

3.240 Å, 0.0000 0.0000 1.0000 c*

Worst diffraction limit after cut-off: 5.837 Å at reflection 20 3 0, in direction 0.989 a*+0.148 b*

Best diffraction limit after cut-off: 3.623 Å at reflection 7 3 41, in direction 0.168 a*+0.072 b*+0.983 c*

Beq: 366.94 [= equivalent overall isotropic B factor on Fs.]

B11 B22 B33

Delta-B tensor: 100.25 100.25-200.49

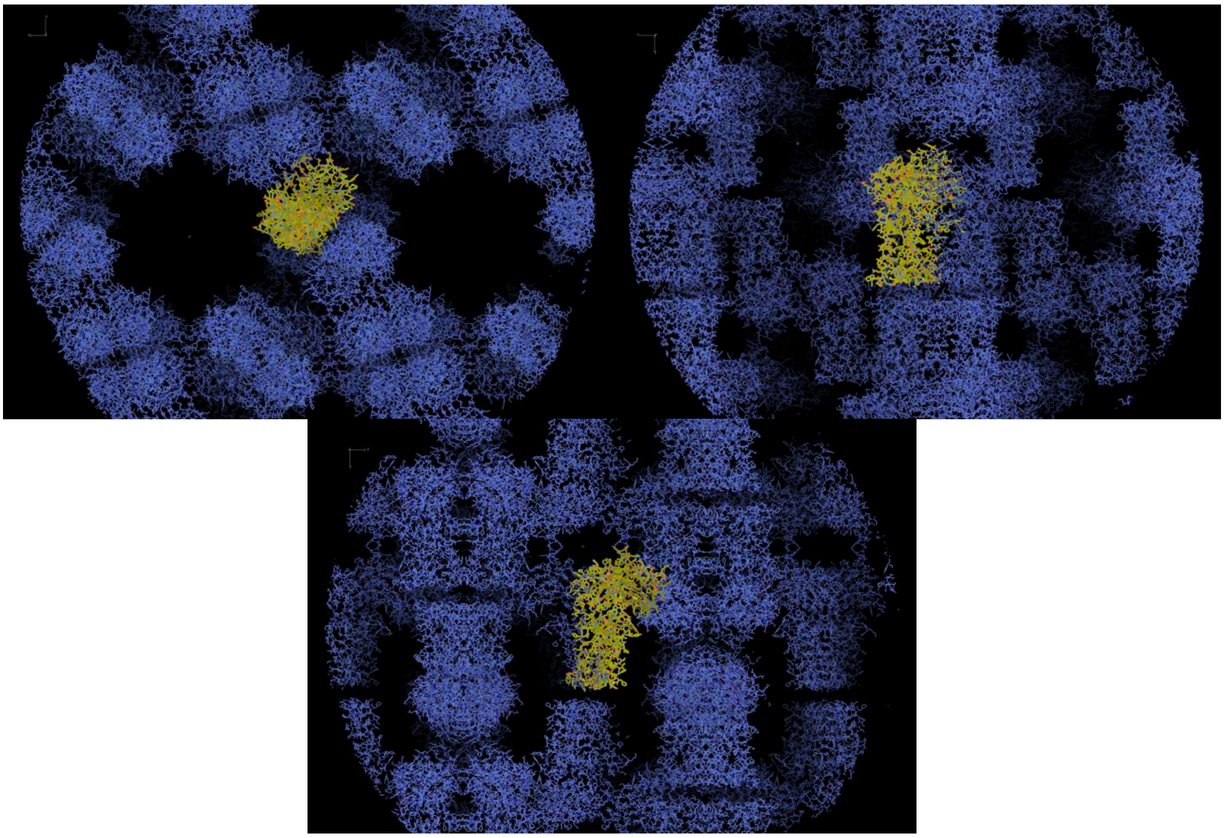

**Fig.S9: Packing in crystal of SpNOX.** Plane X-Y (top left), Plane X-Z (top right), Plane Y-Z (bottom). One asymmetric unit is seen in yellow sticks

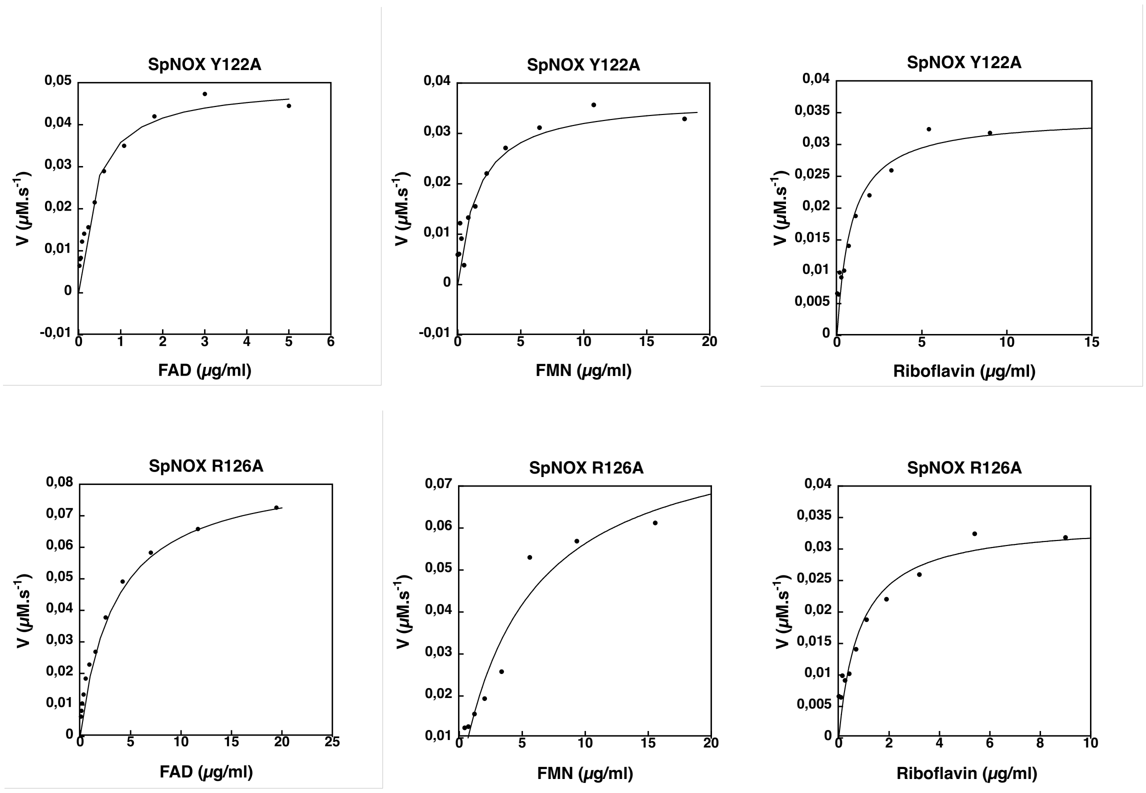


**Fig. S10: Michaelis Menten analysis of SpNOX and SpNOX Y122A and SpNOX R126A as a function of the flavin substrate**. Cytochrome *c* reductase activity was monitored at 550 nm. For all substrates the Michaelis Menten plots are presented.

| | $k_{cat}$ (s-1) | | |
|---|---|---|---|
| | **SpNOX WT** | **Y122A** | **R126A** |
| FAD | 4,23 ± 0.3 | 1,92 ± 0.1 | 3,5 ± 0.1 |
| FMN | 2,46 ± 0.2 | 1,42 ± 0.1 | 2,5 ± 0.1 |
| Riboflavin | 3,23 ± 0.1 | 1,31 ± 0.1 | 1,85 ± 0.1 |

**Table S2:** $k_{cat}$ determined as a function of the mutation in the D-loop and the flavin used. These parameters were extracted from data of Figure S10.

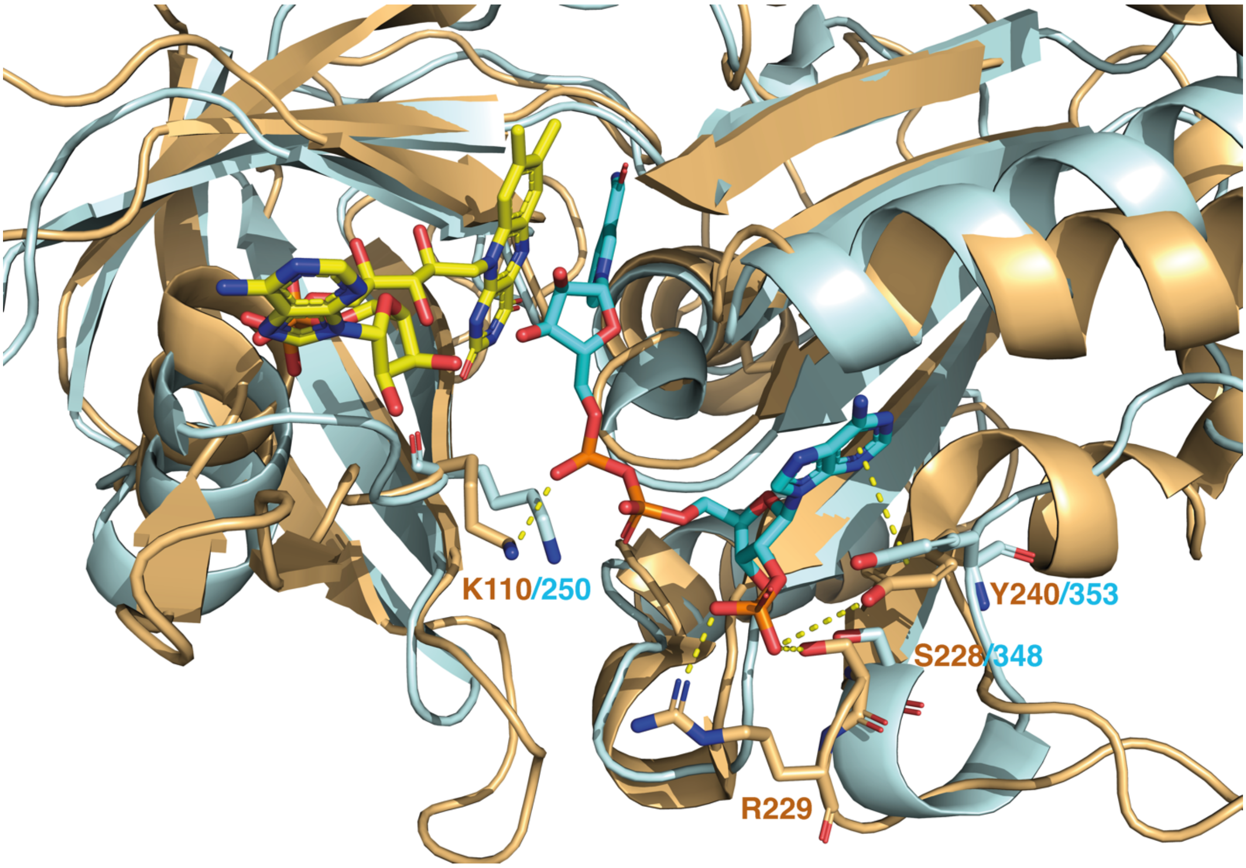


**Figure S11: Overlay of the pea FNR (sand) and the DH domain of SpNOX (pale cyan).** Lysine residues, K110/250, from the FBD domain involved in the pyrophosphate interaction of NADPH are conserved as well as the tyrosine residues, Y240/353, involved in a stacking interaction with the adenine ring and the serine residues, S228/348, binding to the 2'phosphate of the AMP moiety. On the contrary the R229 residue, from pea FNR, interacting with this phosphate is not conserved, in line with the absence of substrate selectivity of SpNOX.

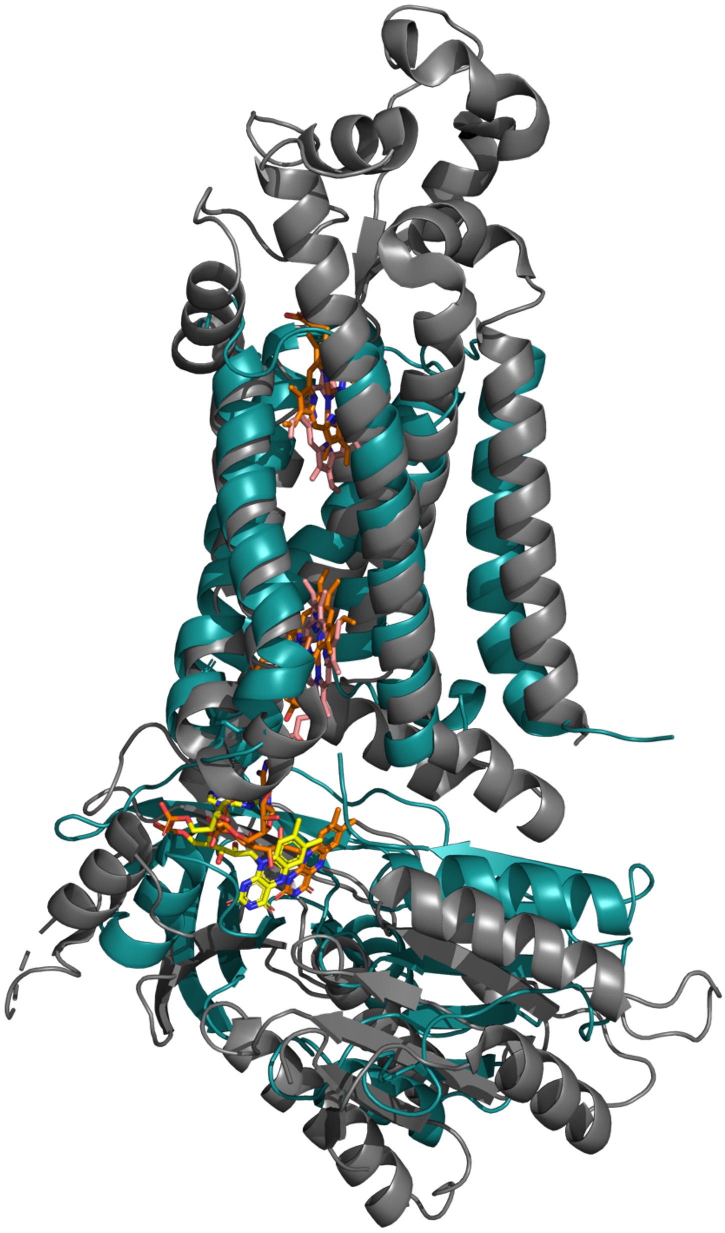

**Figure S12: Superposition of NOX2 and SpNOX.** NOX2 (PDB: 8gz3) is in grey ribbon and SpNOX in blue. FAD is in yellow (NOX2) or orange (SpNOX) sticks; hemes are pink (NOX2) or orange (SpNOX) sticks.
